# Exploring congruent diversification histories with flexibility and parsimony

**DOI:** 10.1101/2023.07.26.550618

**Authors:** Jérémy Andréoletti, Hélène Morlon

## Abstract

1. Using phylogenies of present-day species to estimate diversification rate trajectories – speciation and extinction rates over time – is a challenging task due to non-identifiability issues. Given a phylogeny, there exists an infinite set of trajectories that result in the same likelihood; this set has been coined a congruence class. Previous work has developed approaches for sampling trajectories within a given congruence class, with the aim to assess the extent to which congruent scenarios can vary from one another. Based on this sampling approach, it has been suggested that rapid changes in speciation or extinction rates are conserved across the class. Reaching such conclusions requires to sample the broadest possible set of distinct trajectories.
2. We introduce a new method for exploring congruence classes, that we implement in the R package CRABS. Whereas existing methods constrain either the speciation rate or the extinction rate trajectory, ours provides more flexibility by sampling congruent speciation and extinction rate trajectories simultaneously. This allows covering a more representative set of distinct diversification rate trajectories. We also implement a filtering step that allows selecting the most parsimonious trajectories within a class.
3. We demonstrate the utility of our new sampling strategy using a simulated scenario. Next, we apply our approach to the study of mammalian diversification history. We show that rapid changes in speciation and extinction rates need not be conserved across a congruence class, but that selecting the most parsimonious trajectories shrinks the class to concordant scenarios.
4. Our approach opens new avenues both to truly explore the myriad of potential diversification histories consistent with a given phylogeny, embracing the uncertainty inherent to phylogenetic diversification models, and to select among these different histories. This should help refining our inference of diversification trajectories from extant data.

## 1 Introduction

The study of diversification rates, or the rates at which species arise and go extinct over time, has long been a central topic in evolutionary biology. Early approaches for inferring diversification rates relied on simple heuristic methods, based primarily on observed species ranges in the fossil record (Raup, 1978; Foote, 1994; Jablonski & Chaloner, 1994), and have gradually been expanded to fit into more robust statistical frameworks (Foote, 2000; Alroy, 2014; Silvestro et al., 2014). Another major innovation was to incorporate phylogenetic information from molecular data (Nee et al., 1994) and to model diversification as a continuous branching process of speciation, extinction and sampling (Stadler, 2010). These mathematical developments paved the way for a series of likelihood-based approaches to estimating diversification rates with increasingly complex models, taking into account both rate variation over time (Morlon et al., 2011; Stadler, 2011) and rate heterogeneity across lineages (Alfaro et al., 2009; Rabosky, 2014; Maliet et al., 2019; Barido-Sottani et al., 2020).

However, in recent years, a fundamental problem in statistical inference has been brought to the forefront of diversification rate estimation: the lack of parameter identifiability. For some birth-death diversification models, it is not possible to uniquely determine the underlying parameters of the diversification process from observed data. This issue had already been raised by Kubo & Iwasa (1995), Stadler (2009) and Lambert & Stadler (2013), but Louca & Pennell (2020) demonstrated in full generality that, for a homogeneous birth-death process with speciation and extinction rates that vary freely in time, an infinite combination of speciation and extinction rate curves (usually noted *λ*(*t*) and *μ*(*t*) respectively) yield the exact same likelihood for a given phylogeny of extant species, even in the limit of infinite data. They refer to the infinite set of trajectories (*λ*(*t*), *μ*(*t*)) that are strictly indistinguishable from the phylogeny alone as a “congruence class”. Louca et al. (2021) later extended this non-identifiability result to birth-death processes that also include a fossil sampling rate.

Three main avenues have been followed to address this finding. First, Louca & Pennell (2020) proposed to restrict inferences to identifiable reparametrizations of the model (see also Stadler (2013)), though these “pulled rates” can be challenging to interpret (Helmstetter et al., 2022). The second approach is to work with hypotheses-driven models that are not as flexible as those used by Louca & Pennell (2020), or to add regularity constraints (e.g. priors in a Bayesian inference context) that make the process identifiable (Morlon et al., 2022). Many models that are not as flexible as those used by Louca & Pennell (2020) are indeed identifiable (Legried & Terhorst, 2022a,b). The final option is to embrace the non-identifiability results and to develop methods to explore scenarios within a given congruence class, in order to assess which temporal patterns are conserved across the congruence class and which are not (Höhna et al., 2022; Kopperud et al., 2023). The latter avenue is the one we investigate here, and we link it to the second approach.

Current approaches for exploring congruence classes are based on results from Louca & Pennell (2020), which showed that each congruence class can be uniquely defined by the speciation rate at present *λ*_0_ and a curve called the “pulled diversification rate” curve, defined as 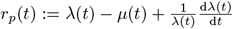 . Given *λ*_0_ and *r*_*p*_, one can explore trajectories in the congruence class by choosing any trajectory for either the speciation rate or the extinction rate and finding the trajectory for the other rate using this equation. In practice, this can be done with a function called `congruent_hbds_model` from the R package *castor* (Louca & Doebeli, 2018). Höhna et al. (2022) developed this approach further with two methods for sampling nearly congruent trajectories based on the following linear approximation (for the time step *t*_*i*_ at iteration *i*): 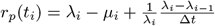. For simplicity we will refer to “congruent trajectories” in what follows, although it should be kept in mind that the trajectories are not exactly congruent, given the linear approximation. The first method is similar to `congruent_hbds_model`, as it involves choosing a trajectory for either the speciation or the extinction rate, and then computing the other trajectory. The second method consists in sampling *λ*(*t*) or *μ*(*t*) in a series of specified function classes, e.g. constant extinction rates with various initial *μ*_0_ values (*λ*_0_ is invariant in each congruence class), Gaussian or Horseshoe Markov random fields (see Magee et al. (2020)), or even a purely uniform sampling at each time step. This aims at truly sampling scenarios in the congruence class, as opposed to hand-picking few of them. The authors implemented these approaches in a new R package called CRABS (Congruent Rate Analyses in Birth-Death Scenarios). In a follow-up paper, Kopperud et al. (2023) used CRABS to investigate the speciation curves resulting from sampling extinction curves, as sampling speciation curves to recover congruent extinction curves is trickier (see below). With this approach, the authors showed that moderate rate changes are not necessarily recovered across congruent models, but that sharp shifts often are.

One main limitation of these approaches is that they strongly constrain the rate trajectory that is fixed or sampled (*λ* or *μ*), as opposed to the trajectory that is recovered through the congruence class, hence limiting the variability of patterns that can be observed. Even the CRABS option of choosing a series of uniformly sampled points imposes the absence of any recurrent pattern on the rate trajectory that is sampled. As we illustrate in the paper, this can be problematic. First, if only one strategy is used (for example sampling mostly extinction rates as in Kopperud et al. (2023)), we may misleadingly conclude that the inference is robust across the congruence class. If instead several alternative sampling strategies are used, as recommended by Höhna et al. (2022), we may find very distinct congruent scenarios, some of them not very plausible biologically, with difficulties to reach conclusions about the true diversification dynamics. Second, these sampling schemes seem to have difficulties sampling diversification histories close to the one used to define the congruence class. We certainly want to be able to sample different congruent trajectories, but recovery of the initial trajectories should be a prerequisite.

In this paper, we develop alternative sampling strategies of the congruence class, as well as a potential additional filtering step which consists in selecting the most parsimonious trajectories. We implement these approaches in CRABS. Finally, we illustrate their utility by applying them to a simulated scenario as well as mammals’ diversification.

## 2 Methods

### 2.1 Sampling on the congruence curve

Given a congruence class uniquely characterized by the pulled diversification rate trajectory *r*_*p*_ and the speciation rate at present *λ*_0_, we aim to simultaneously sample speciation (*λ*) and extinction (*μ*) rate trajectories within the class, instead of constraining the functional space of one of the two trajectories. Following Höhna et al. (2022), we discretize time into small steps of size ∆*t* and use the linear approximation of *r*_*p*_ mentioned in the Introduction. For each iteration *i*, we define the *congruence curve* as the set of (*λ*_*i*_, *μ*_*i*_) pairs such that the resulting trajectories remain within the congruence class. Rearranging the linear equation for *r*_*p*_, the congruence curve has the following form (see Figure 1):

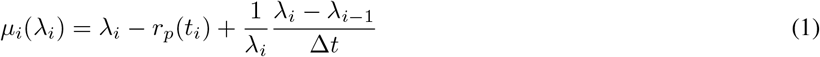

At present *t*_0_, *λ*_0_ is given and we sample *μ*_0_ in a given distribution (see details below). Next, going backwards in time, we sample *λ*_*i*_ and *μ*_*i*_ simultaneously along the congruence curve at each iteration *i*. This allows for a more flexible and balanced sampling within the congruence class.

**Figure 1.**
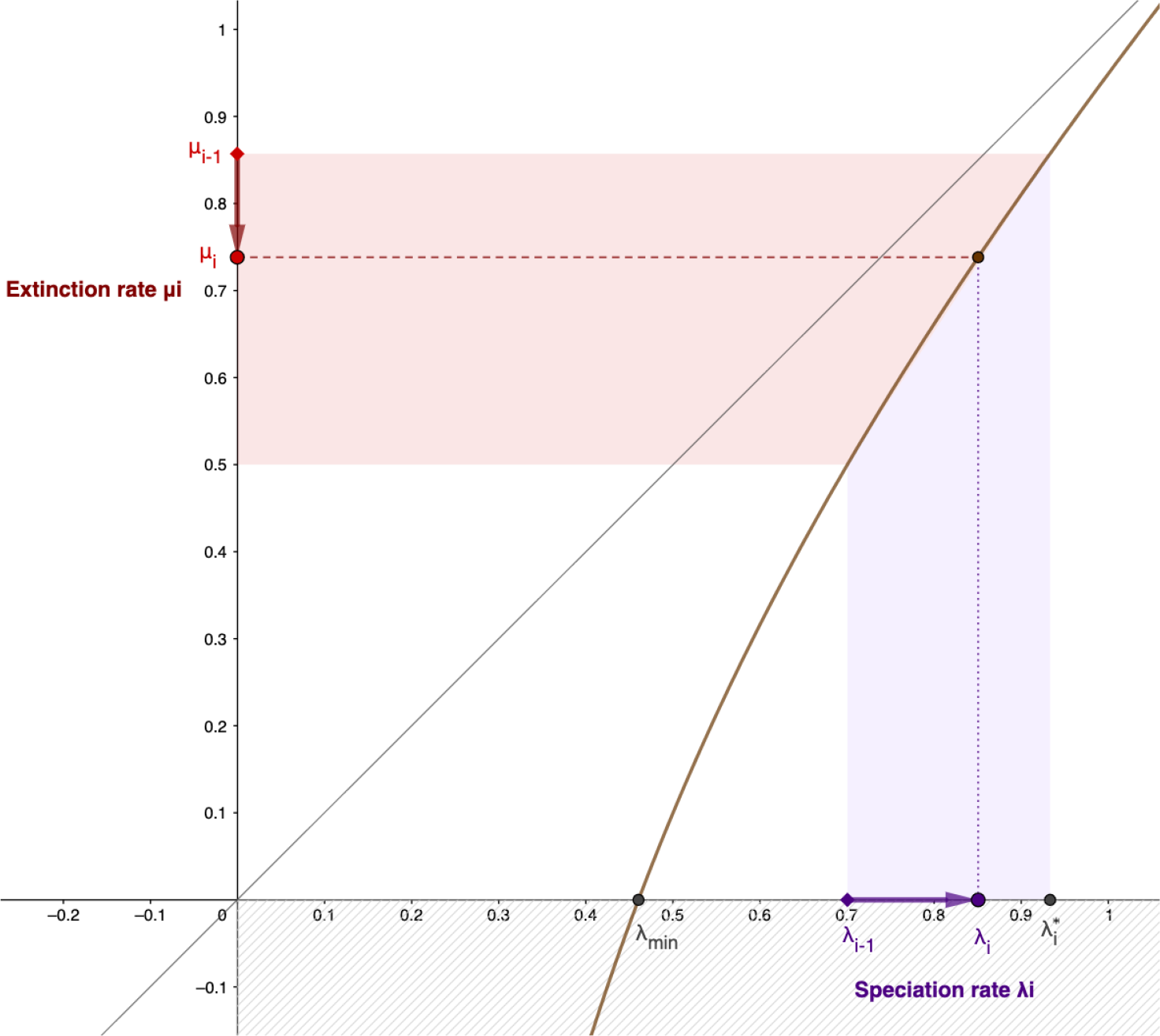
Sampling the congruence curve. Space of parameters (*λ*_*i*_, *μ*_*i*_) that can be sampled at a given iteration *i*. At each step, *λ*_*i*_− _1_ and *r*_*p*_(*t*_*i*_) define a specific congruence curve (in brown) along which (*λ*_*i*_, *μ*_*i*_) can be sampled so that the trajectory remains in the congruence class ; the fact that it is below the 1:1 gray line in this illustration indicates a positive net diversification trend. We further impose the constraint of a minimal *λ*_*min*_ value to avoid negative extinction rates (dashed area). Note that choosing *λ*_*i*_ = *λ*_*i*_ −_1_ implies a constant speciation rate while choosing 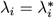 implies a constant extinction rate ; we will be sampling a position on the curve relative to these two values. The figure represents the situation when 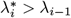, in which case *λ*_*i*_ *> λ*_*i*_ −_1_ and *μ*_*i*_ *< μ*_*i*−1_, but we can also later have 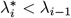, in which case *λ*_*i*_ *< λ*_*i*−1_ and *μ*_*i*_ *> μ*_*i*−1_.

Before sampling on the congruence curve, we compute two useful quantities : 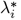, the *λ*_*i*_ value such as *μ*_*i*_ = *μ*_*i*−1_, and 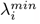, the minimum *λ*_*i*_ value that maintains non-negative *μ*_*i*_.

Assuming that ∀*i, λ*_*i*_ *>* 0:

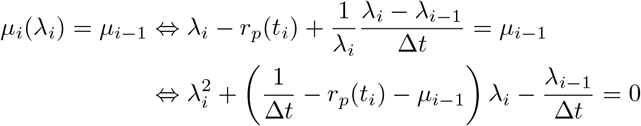

The only positive solution of this quadratic polynomial (given that the congruence curve is monotonic for *λ*_*i*_ *>* 0) is the following:

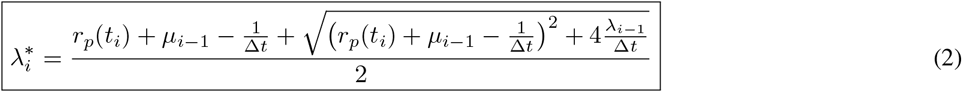

From this equation, we can directly deduce 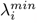, the minimum *λ*_*i*_ value that keeps *μ*_*i*_ non-negative, by setting *μ*_*i*−1_ to 0:

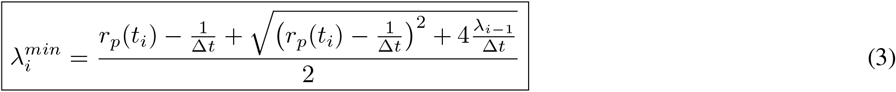

The iterative sampling along the congruence curve is illustrated in Figure 1, plotted with Geogebra (Hohenwarter, 2023).

### 2.2 Sampling approaches

At a given iteration *i*, on one extreme *λ* is set constant (*λ*_*i*_ = *λ*_*i*−1_), implying that the time-varying pattern in the congruence class will be reflected in *μ*_*i*_. On the other extreme *μ* is set constant 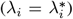 and *λ*_*i*_ conveys all the information. In order to obtain other sampling approaches, we select a position *p*_*i*_ relative to *λ*_*i*−1_ and 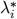 . The various proposed approaches make different choices for selecting *p*_*i*_. Below, “*p*-trajectory” refers to the trajectory of *p*_*i*_ as we move through time steps *t*_*i*_ from the present to the past.

#### 2.2.1 Fixed position on the congruence curve: Constant *p*-trajectory

Our first approach consists in taking at each iteration *i* the same relative position *p*_0_ between *λ*_*i*−1_ and 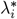 (i.e. *p*_*i*_ is constant). Here is the complete procedure:

1. Choice of the position : *p*_0_ ∼ *Beta*(*β*_1_, *β*_2_)
2. Initialization : 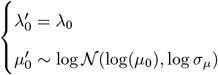
3. For each iteration *i* ≥ 1
  a. Compute 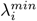and 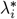
  b. Take 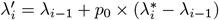, i.e. at a relative position *p*_0_ between *λ*_*i*−1_ and 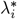
  c. If 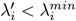, set 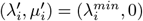 to avoid negative *μ* values
  d. Else, compute 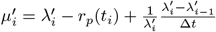 according to the congruence relationship

Note that *p*_0_ = 0 implies a constant speciation rate throughout a clade’s history (barring negative extinction rates), while *p*_0_ = 1 implies a constant extinction rate. Our method explores intermediate trajectories between these two poles. As a consequence, even if the algorithm above only samples *λ* before retrieving *μ* through the congruence curve, *μ* is still included in the procedure through the computation of 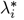 . Note also that at each step this procedure forces either speciation rates to increase and extinction rates to decrease (if 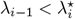), or the reverse (if 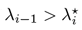), which is in itself a constraint favoring parsimonious trajectories (as the data could typically also be explained by arbitrary large simultaneous increases in both rates).

The parameters of the Beta distribution (*β*_1_, *β*_2_) can be set to (1, 1) to sample uniformly between 0 and 1, or changed to different values in order to get more samples of specific trajectories (e.g. nearly constant lambda or mu, and all intermediates).

#### 2.2.2 Autocorrelated position on the congruence curve: Markov Random Field (MRF) *p*-trajectory

This second approach is very similar to the previous one, with the distinction that *p*_*i*_ is not constant at each iteration *i*, but instead follows an autocorrelated process starting at *p*_0_. Following Magee et al. (2020), we use either a Gaussian Markov random field (GMRF), a model in which *p*_*i*_|(*p*_*i*−1_, *γ*) ∼ 𝒩 (*p*_*i*−1_, *γ*^2^), or a Horseshoe Markov random field (HSMRF), a model in which *p*_*i*_|(*p*_*i*−1_, *ζ*) ∼ *ℋorseshoe*(*p*_*i*−1_, *γ*^2^). *γ* is a global scale parameter that controls the smoothness of the overall field (shrinking all jumps towards zero when *γ* tends towards zero). In the case of the HSMRF, the fat-tailed nature of the Horseshoe distribution allows for rare occurrences of larger jumps in the trajectory.

1. Drawing the *p*-trajectory
  a. **Global scale parameter** controlling the degree of regularization (i.e. steering towards “simpler” trajectories) : *γ* ∼ *halfCauchy*(0, *ζ*) This parameter is itself controlled hierarchically by the hyperparameter *ζ*, which can be obtained with the CRABS function *get*.*gmrf*.*global*.*scale* such that the expected number of 2-fold *p* changes given the number of time steps equals log(2). It can be further adjusted manually with a scaling factor *σ*_MRF_.
  b. Trajectory: starting value *p*_0_ ∼ *Beta*(*β*_1_, *β*_2_), then

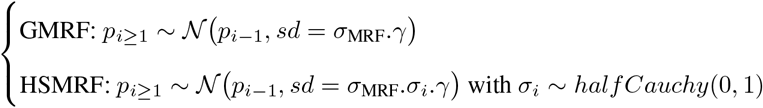

*σ*_*i*_ is a **local shrinkage parameter** allowing for rare larger jumps.
  c. Centering: by default, such autocorrelated trajectories will see their variance increase over the time steps, resulting in more variable dynamics in the past than in the present. To cancel this effect, the *p*-trajectory is centered around the *p*_*i*_ value at a randomly sampled time step.
  d. Rescaling: here the *p*-trajectories are not necessarily bounded between 0 and 1, so it is possible for *λ*_*i*_ and *μ*_*i*_ to both increase (*p*_*max*_ *>* 1) or decrease (*p*_*min*_ *<* 0) simultaneously. We however rescale the *p*-trajectories whenever they leave predefined bounds. We used *p*_*min*_ = −0.05 and *p*_*max*_ = 1.05, which worked well in practice, and we implemented them as default values.
2. Initialization : 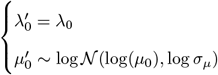
3. For each iteration *i* ≥ 1: same as before with *p*_*i*_ instead of *p*_0_

#### 2.2.3 MRF *p*-trajectory with a global regularity filter

It is possible to construct congruent diversification rate trajectories with arbitrarily large and sudden rate changes, however these diversification histories may be less plausible biologically than smoother ones. If one follows the parsimony principle, simple trajectories that explain the observed data as well as more complex ones should be preferred Morlon et al. (2022). This *a priori* intuition can be incorporated by generating many congruent diversification trajectories, for example following the MRF *p* sampling approach, and selecting those that do not exhibit such sudden sharp changes. In practice, selecting “smooth” or “regular” rate trajectories can be done by ranking them based on a quantitative measure of their smoothness, and disregarding those that rank poorly. We tested three quantitative measures of the regularity/smoothness of diversification rate trajectories:

- The *ℒ* _1_ norm on all the step-wise variations in both speciation (*λ*_*i*_) and extinction rates (*μ*_*i*_), for *i* ∈ [[0; *N* ], where *N* is the number of time steps :

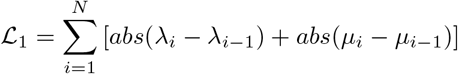

The *ℒ* _1_ norm sums the absolute differences between consecutive speciation and extinction rates. Rate trajectories with low *ℒ* _1_ norms tend to have few and small rate variations.
- The *ℒ* _2_ norm on step-wise variations:

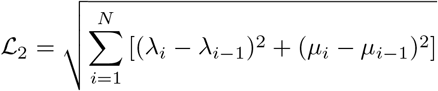

The *ℒ* _2_ norm, also known as the Euclidean norm, squares step-wise differences before summing and taking the square root. This has the effect of penalizing larger jumps more severely than smaller ones.
- The second-order *ℒ* _1_ norm on derivative shifts in both speciation and extinction rates:

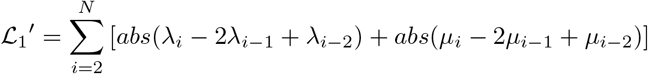

The second-order *ℒ* _1_ norm 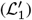 focuses on the change in the rate of change, effectively measuring the volatility in the trajectory. This offers an additional layer of smoothness by considering how rate changes are themselves changing.

These norms serve as a criterion for selecting parsimonious diversification scenarios. After generating many congruent diversification trajectories, we rank them according to the value of the selected norm. We can then decide, for instance, to exclude the 10% most “irregular” trajectories, i.e. the 10% with highest values of the norm. The number of initial samples must be high enough to ensure that enough trajectories are retained. We recommend generating at least 100 times the number of retained samples. This approach aligns well with the principle of parsimony: among models that explain the data equally well (congruent trajectories have the same likelihood), a choice is made to favor simpler models, i.e. smoother trajectories.

#### 2.2.4 Implementation in CRABS

We implemented our new sampling methods in R as extended functions of the CRABS package (R Core Team, 2018; Höhna et al., 2022). As in CRABS, two functions are called. First, the function `sample.basic.models.joint` (modified from `sample.basic.models`), which in its default setting returns a sampler for HSMRF *p* -trajectories, and can return a sampler for constant *p*-trajectories by choosing a zero scaling factor (`mrf.sd.scale = 0`). We chose the HSMRF strategy as a default, since the GMRF is less flexible. Second, the function `sample.congruence.class` returns the samples; we simply added the argument `sample.joint.rates` to this function, and the user should specify to use the option `rate.type = “joint”`.

We also implemented the function `full.plot.regularity.thresholds` that ranks the sampled trajectories according to their regularity values, selects the *x*% most regular, and displays the corresponding trajectories. The arguments `filtering_fractions` (by default 1%, 5%, 20% and 90%), `penalty` (among “L1”, “L2” and “L1_derivative” for the derivative shifts regularity criterion) and `rates` (vector including “lambda” for speciation, “mu” for extinction, “delta” for net-diversification, “epsilon” for turnover) give control over the specific filtering being performed.

### 2.3 Illustrations

#### 2.3.1 Simulated scenario

In order to illustrate the utility of our sampling approach in comparison with previous approaches, we analyse a mock reference scenario with a declining speciation rate 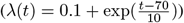 and a mostly constant extinction rate with a sudden spike 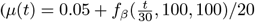 with *f*_*β*_ the probability density function of the Beta distribution). This model corresponds to an often-observed scenario of early-burst diversification (Moen & Morlon, 2014) combined with a mass extinction event. We first apply the previous CRABS functions (Höhna et al., 2022), sampling 20 congruent HSMRF trajectories of either speciation or extinction rates using the default options, and summarize relative rate changes over time with the recommended threshold of 0.02 (in unit of rate per time) for significant increases and decreases.

We then carry out our two new procedures on the same scenario, sampling 20 constant *p*-trajectories and 20 HSMRF *p*-trajectories, each corresponding to a joint sampling of speciation and extinction rates.

#### 2.3.2 Mammalian diversification

We analyse the diversification dynamics of mammals, using the dated mammalian phylogeny constructed by Álvarez-Carretero et al. (2022), which integrates information from 72 mammal genomes. More specifically, we used the subset of this timetree that excludes subspecies (634 among the 4705 initial lineages), as used in Quintero et al. (2022).

We conducted an initial diversification analysis in RevBayes (Höhna et al., 2016) assuming auto-correlated piecewise-constant speciation and extinction rates trajectories described by a HSMRF prior distribution (see Magee et al. (2020) and the corresponding RevBayes tutorial^1^). We used a single sampling fraction (0.7287) and 30 time intervals whose limits were determined during the inference process. We visualized the results using the RevGadgets R library (Tribble et al., 2022; R Core Team, 2018).

Next, we generated 5000 joint HSMRF *p*-trajectories on the congruence curve, using 500 time points (∆*t* ≈ 400, 000 years), and filtered them based on their regularity values, using each of the three available norms. For illustration purposes, we sampled four sets of 50 trajectories within those that fell below the 1st, 5th, 20th, and 90th centiles of regularity values.

Finally, since using a prior distribution on rate variations tends to smooth diversification trajectories, we carried a similar analysis without such prior. We conducted a diversification analysis using piecewise-constant rate trajectories with independent estimates for each episode and only 10 time intervals. Next, we constructed congruent curves and filtered them based on their regularity values, as detailed above.

## 3 Results

### 3.1 Simulated Scenario

Applied to the mock scenario with a decline in speciation rates and a spike in extinction, the methods of Höhna et al. (2022), which sample either speciation or extinction rate trajectories, lead to two possibilities depending on whether one or the other is sampled (Fig. 2). Sampling the extinction rate trajectory allows recovering the decline in speciation rate, but also infers a systematic short increase in the speciation rate at the time of the (unaccounted for) extinction peak (Fig 2 a,c). Sampling the speciation rate trajectory instead allows recovering the extinction peak, but also infers a spurious initial increase in extinction rate reflecting the (unaccounted for) decline in speciation (Fig 2 b,d). In addition, all inferred extinction rates at the origin of the clade are negative, illustrating the difficulty of sampling the congruence class through speciation trajectories.

**Figure 2.**
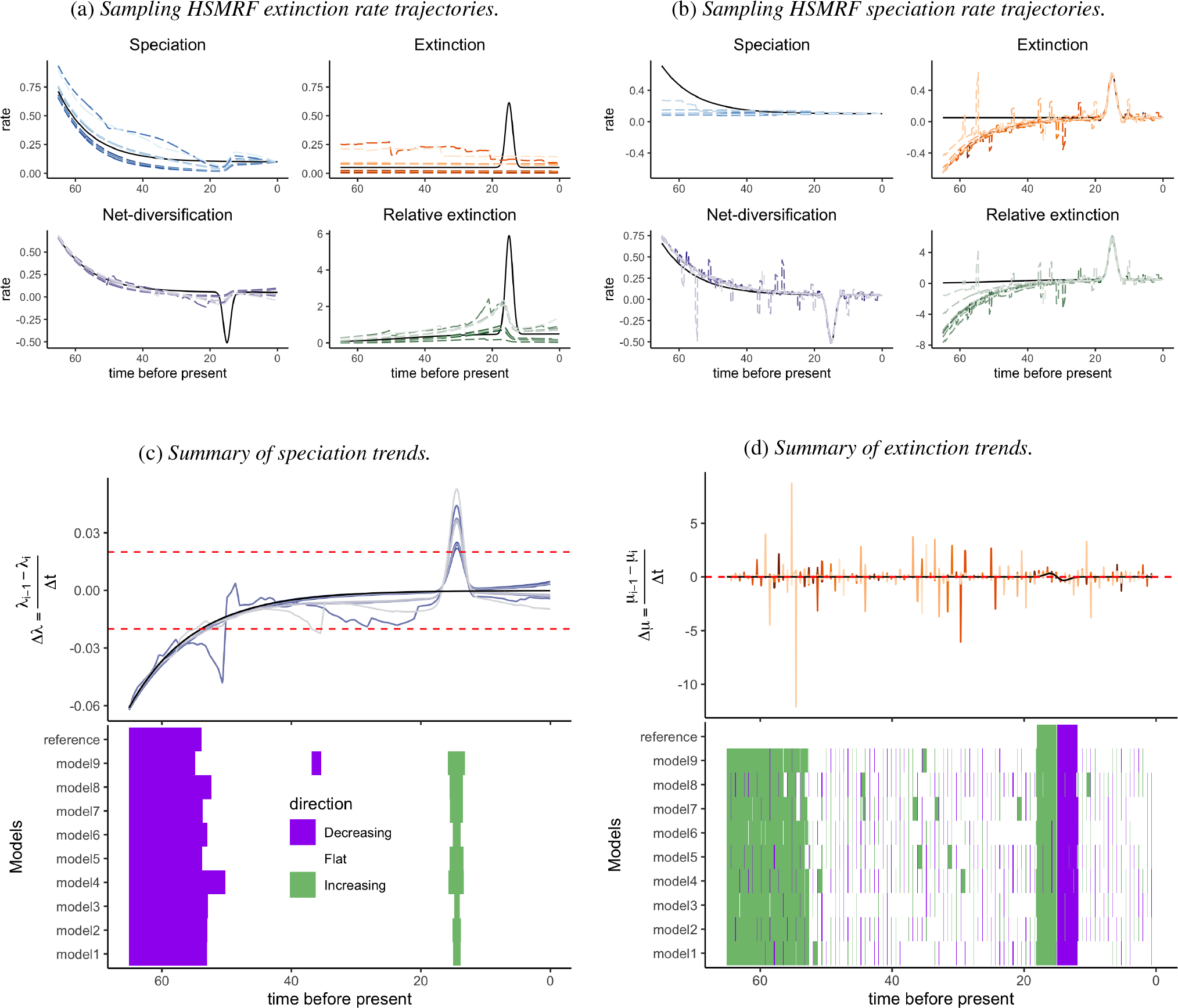
Illustration of current limitations in the exploration of congruence classes. (a,b) Rate trajectories over time for the reference scenario (black line) and 20 congruent scenarios (dashed lines) obtained with previous CRABS functions (Höhna et al., 2022). In (a), 20 extinction rate trajectories are first sampled using the HSMRF process, and corresponding congruent speciation rate trajectories are computed for each. In (b), 20 speciation rate trajectories are first sampled using the HSMRF process, and corresponding congruent extinction rate trajectories are computed for each. (c,d) Top: relative rate changes over time, with red dashed lines demarcating the 0.02 significance threshold. Bottom: summary across 10 congruent models with increases in green and decreases in purple; the first row at the top corresponds to the reference scenario. Note that in both cases, no sampled congruent trajectory shows patterns close to the reference scenario used to construct the congruence class, and that the macroevolutionary signal is reflected either entirely on speciation rate, or entirely on extinction rate.

In contrast, the sampling approaches we developed were able to sample congruent scenarios with improved flexibility, including around the initial model (Fig. 3). With the constant *p*-trajectory sampling approach (Fig. 3a), extinction rates ranged from completely flat to the large initial spike, and speciation rates showed a steady decline, with sometimes a small decrease at the time of the sharp decline in the net diversification rate, corresponding to the cases when the spike is not fully reproduced in the extinction curve. HSMRF *p*-trajectories (Fig. 3b) result in similar sampled trajectories with more variability, including congruent scenarios with higher speciation and extinction rates in the past.

**Figure 3.**
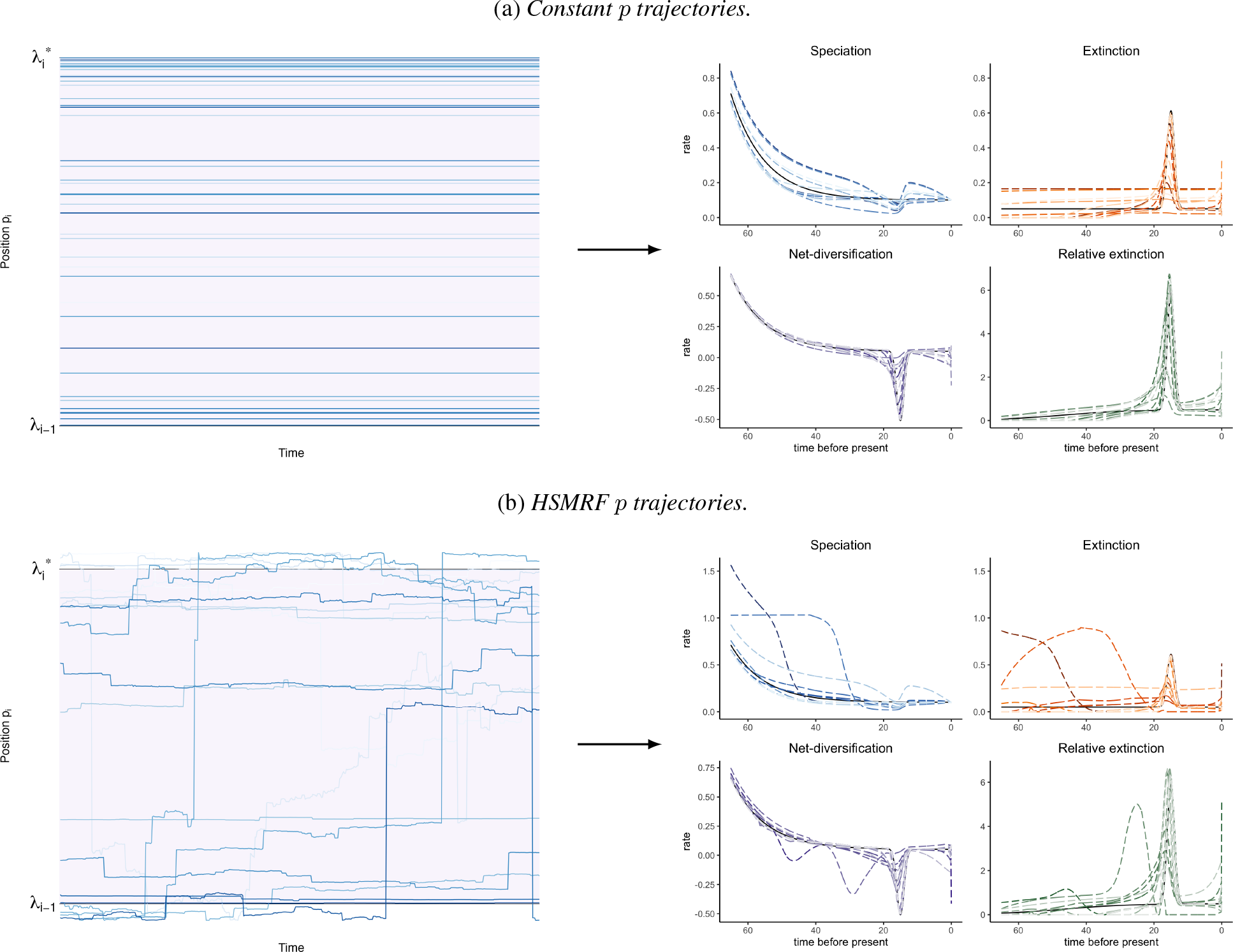
Two approaches to jointly sample speciation rate and extinction rate trajectories. Each blue *p*-trajectory on the left panel corresponds to rate trajectories on the right panel. The black line represents the mock diversification scenario used in Figure 2, other trajectories (dashed lines) are sampled from its congruence class. Their trends are summarized in Sup. Fig. A.3. (a) Evolutionary rates are sampled through time always at the same position between between *λ*_*i*_ − _1_ (*λ* as constant as possible without negative extinction) and 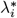(constant *μ*). (b) Evolutionary rates are sampled through time similarly, except that the position relative to *λ*_*i*−1_ and 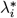 follows a HSMRF process.

### 3.2 Mammalian diversification histories

The diversification analyses on the mammalian phylogeny assuming auto-correlated piecewise-constant speciation and extinction rates trajectories in RevBayes suggest a 10-fold jump in speciation rates (from 0.01 to 0.1 *Msy*^−1^) some 80-85 million years ago and a constant, low extinction rate (Figure 4). Congruent mammalian diversification histories that are selected under the most stringent regularity constraints (the 1% most regular trajectories) match closely the initial trajectories (Figure 4).

**Figure 4.**
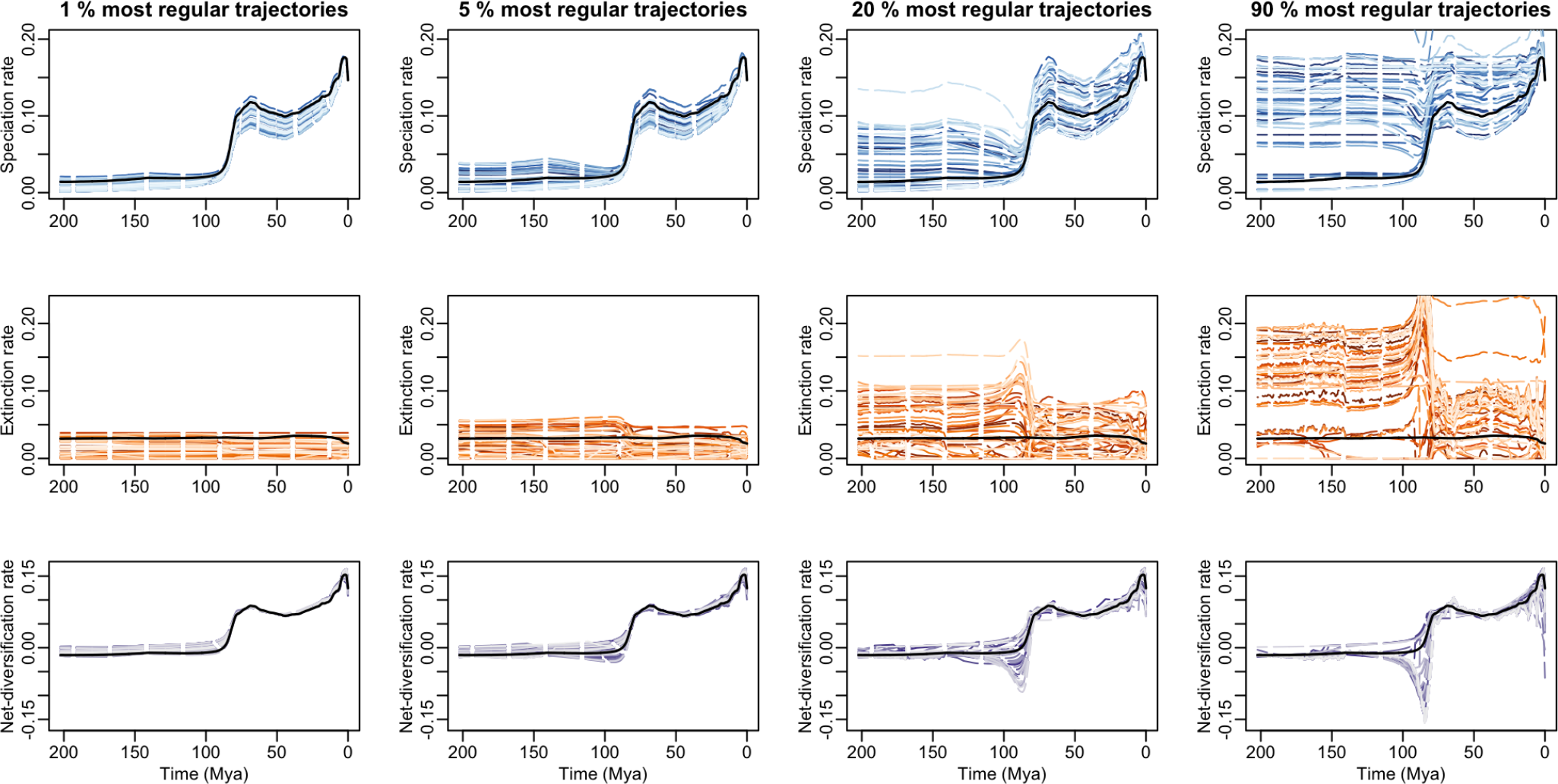
Congruent mammalian diversification trajectories for increasing regularity thresholds. The median rate trajectory inferred in RevBayes is displayed in black, together with congruent speciation (top), extinction (middle) and diversification (bottom) rates in dashed lines. Joint HSRMF *p*-trajectories were generated on the congruence curve, and selected based on their regularity. The most stringent regularity criterion is represented on the left, with 50 trajectories sampled below the first centile of regularity values. Similar plots are provided using the 5th, 20th, and 90th centiles, from left to right.

Some of the more complex trajectories paint a different picture, where the 80-85 Mya signal is driven both by fluctuations in speciation rates and by increased extinction rates before and during this period. These congruent trajectories are also characterised by markedly lower diversification rates during the transition period. Similar patterns are found with other regularity penalties (see Supplementary Figures B.1 and B.2).

When rates are inferred without any regularity prior on piecewise variation, the resulting speciation and extinction trajectories from RevBayes show more variation, as expected (Supplementary Figure B.3). In this case, the trajectories that exhibit the highest regularity within the congruence class bear a closer resemblance to those generated using the regularizing prior, rather than resembling the reference piecewise diversification curve without prior regularization. In particular, the most regular trajectories have almost constant extinction rates (Supplementary Figure B.3).

## 4 Discussion

We introduced a novel approach for sampling alternative diversification scenarios in a given congruence class. By jointly sampling congruent speciation and extinction rate trajectories, our approach provides a more thorough representation of congruence classes than previous methods which sampled entirely one of the two trajectories before deducing the other. Although a congruence class contains very distinct diversification scenarios, which are indistinguishable based on their likelihood to have produced a given extant phylogeny, these scenarios are not equally regular. We implemented the possibility to select congruent scenarios based on several regularity criteria. As we illustrated in our empirical analysis of mammals, this filtering step can shrink the congruence class to a unique diversification scenario. Our developments therefore provide the possibility to explore a wide range of equally likely diversification scenarios while keeping biological interpretability.

### 4.1 Embracing the uncertainty

In the face of the non-identifiability of birth-death diversification models with unconstrained time-variable rates, it has been proposed to restrict inferences to identifiable reparametrizations of the model Louca & Pennell (2020). Unfortunately, such reparametrizations come at the cost of limited biological interpretability (Helmstetter et al., 2022). Another approach is to work with hypothesis-driven models (Morlon et al., 2022). In this case the models often are identifiable, but this might give overconfidence in misleading results, as the conclusions may be highly dependent on the hypotheses being made. These two approaches have the common disadvantage to force us to restrict our analysis to identifiable patterns, rather than to formulate the models we believe best reflect the processes at play regardless of whether they are identifiable or not (Gustafson et al., 2005; Gustafson, 2015). Yet another approach is to rely on a Bayesian framework, in which inference of posterior parameter distributions is always theoretically identifiable. However, when the underlying model is non-identifiable, the final outcomes of Bayesian inference are highly dependent on the choice of priors, reflecting our current beliefs. In such situations, it can be beneficial to embrace the uncertainty and complement the analyses with an exploration of congruent trajectories, which represent a less subjective probing of the potential processes at work. Such exploration can help assess the results’ sensitivity to the *a priori* hypotheses and/or the prior choice: congruent trajectories that deviate significantly from the originally inferred posterior trajectories but appear biologically realistic suggest that other hypotheses or priors may warrant further scrutiny.

Höhna et al. (2022) paved the way to the exploration of congruence classes, yet their sampling approach is not as exhaustive as it may seem. Sampling one rate trajectory arbitrarily as a first step before computing the other trajectory to obtain a congruent scenario in a second step forces signal in the data to be captured in the second trajectory. One risk, given the difficulty of obtaining trajectories with positive extinction rates when sampling speciation rate trajectories, is to sample mostly extinction rate trajectories (see e.g. Kopperud et al. (2023)), and therefore to force signal in the data to be entirely captured in the speciation rate trajectory. This provides a biased exploration of congruence classes that may give an impression of robustness that could fall apart when exploring the class more exhaustively. Sampling congruent speciation and extinction rate trajectories jointly, as we do here, provides an efficient approach for exploring the congruence class while allowing signal in the data to be captured by both speciation and extinction rate curves, therefore truly embracing the uncertainty.

### 4.2 Shrinking the uncertainty

Although embracing the uncertainty by exploring congruence classes provides a fair representation of the diversity of scenarios that cannot be distinguished from their likelihood, it can be useful to shrink this uncertainty using other criteria, such as regularity criteria. Facing equally likely hypotheses is not uncommon in the process of scientific inquiry, but this has never been an insurmountable barrier in the face of Occam’s razor. Indeed, it is always possible to create more complex models to explain any set of observations, but hypotheses with equal likelihoods need not be equally plausible. Among a set of equally likely hypotheses, it is common practice to assign higher prior probabilities to the most parsimonious ones (Morlon et al., 2022). We have implemented several regularity criteria to perform this selection. Interestingly, in our empirical illustration on mammals, the most regular congruent diversification scenarios constructed from two distinct initial piecewise-constant diversification rate models, one implementing a smoothing prior on rate variations and the other one not, match closely the initial scenario obtained with the smoothing prior. This suggests a certain degree of stability in parsimonious congruent diversification rate trajectories constructed from distinct initial diversification models. Additionally, it indicates that currently-used diversification models that penalize rate variation already perform well in selecting the most regular trajectories within congruence classes.

Visualizing congruent rates under varying stringency thresholds (as depicted in Fig. 4) eliminates the need to make an arbitrary choice for the threshold and transparently displays the distribution of possible trajectories, leaving the interpretation to depend on the reader’s willingness to favor the “simplest” histories that can explain the data.

### 4.3 Perspectives

Congruence classes have been characterized beyond classical birth-death models, for example in the context of the fossilized birth-death process. This process contains an additional rate for incorporating samples over time into the phylogeny, interpreted as fossils in macroevolutionary studies and genetic samples in epidemiological analyses (Andréoletti et al., 2022; MacPherson et al., 2022). Although this model has been demonstrated to be non-identifiable in the (macroevolutionarily unrealistic) case where lineages go extinct upon sampling (Louca et al., 2021), a more general demonstration is lacking and a method for sampling within this congruence class has not been established. The methodology presented in this study could potentially be extended for the joint sampling of the three rate parameters.

## 5 Conclusion

Embracing the uncertainty inherent to phylogenetic diversification analyses by exploring congruence classes can help us avoid missing realistic diversification scenarios that cannot be ruled out by the observed data. As we nevertheless strive to narrow down this uncertainty, we contend that restricting ourselves to simpler identifiable quantities that are hard to interpret biologically is not the most promising strategy. Instead, introducing additional independent data types (e.g. fossils, biomarkers of ancient organisms) and prior knowledge while navigating between embracing and shrinking the uncertainty, can provide a reliable and nuanced portrayal of past diversification histories, ultimately enhancing our comprehension of the complex evolutionary processes that shape ecological systems.

## 6 Acknowledgements

We thank Bjørn T. Kopperud, Sebastian Höhna, Andrew F. Magee for their constructive feedback and reviewing the code, and Lucas Rey and Nathanaël Boutillon for discussions on the sampling method.

## 8 Availability of data and materials

Our functions have been integrated into the CRABS R package, available on Github (https://github.com/afmagee/CRABS). The code and data corresponding to the analyses presented in this article are available at https://doi.org/10.5281/zenodo.8386963.

## 9 Funding information

J.A. is supported by a doctoral fellowship from the École Normale Supérieure de Paris. HM acknowledges support from the European Research Council (grant CoG-PANDA).

### 10 Authors’ contributions

Both authors contributed to the design of the project, the discussion of the research, and the writing and editing of the article. Jérémy Andréoletti wrote the code, performed the analysis and wrote the first draft.

## Appendix A Simulated scenario

### A.1 Exponential decrease of speciation rates or exponential increase of extinction rates

**Figure A.1:**
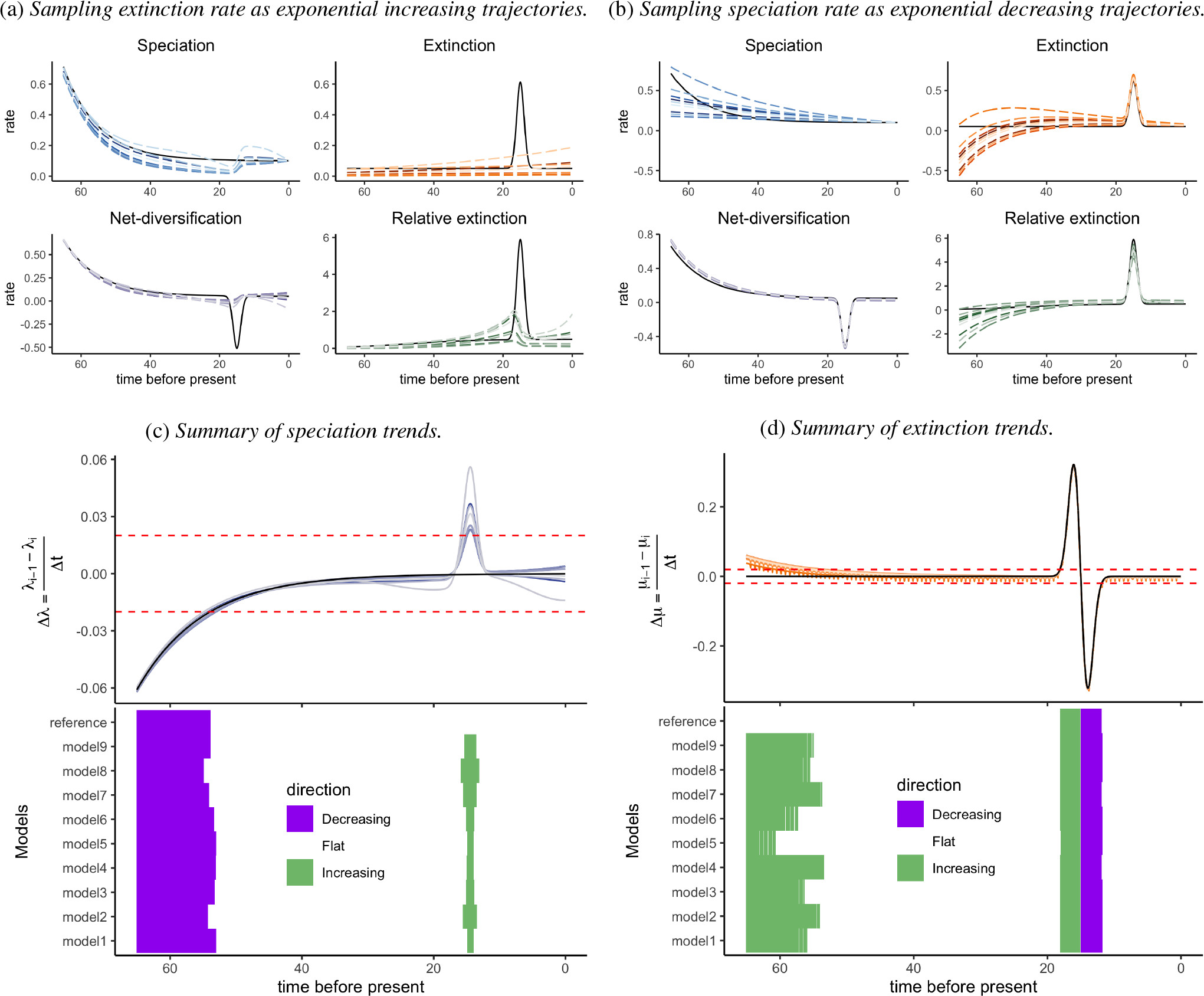
Illustration of diversification scenarios recovered with current congruence class sampling methods. Same as Figure 2, with speciation or extinction rates sampled as exponential trajectories.

### A.2 GMRF trajectories of speciation and extinction rates

**Figure A.2:**
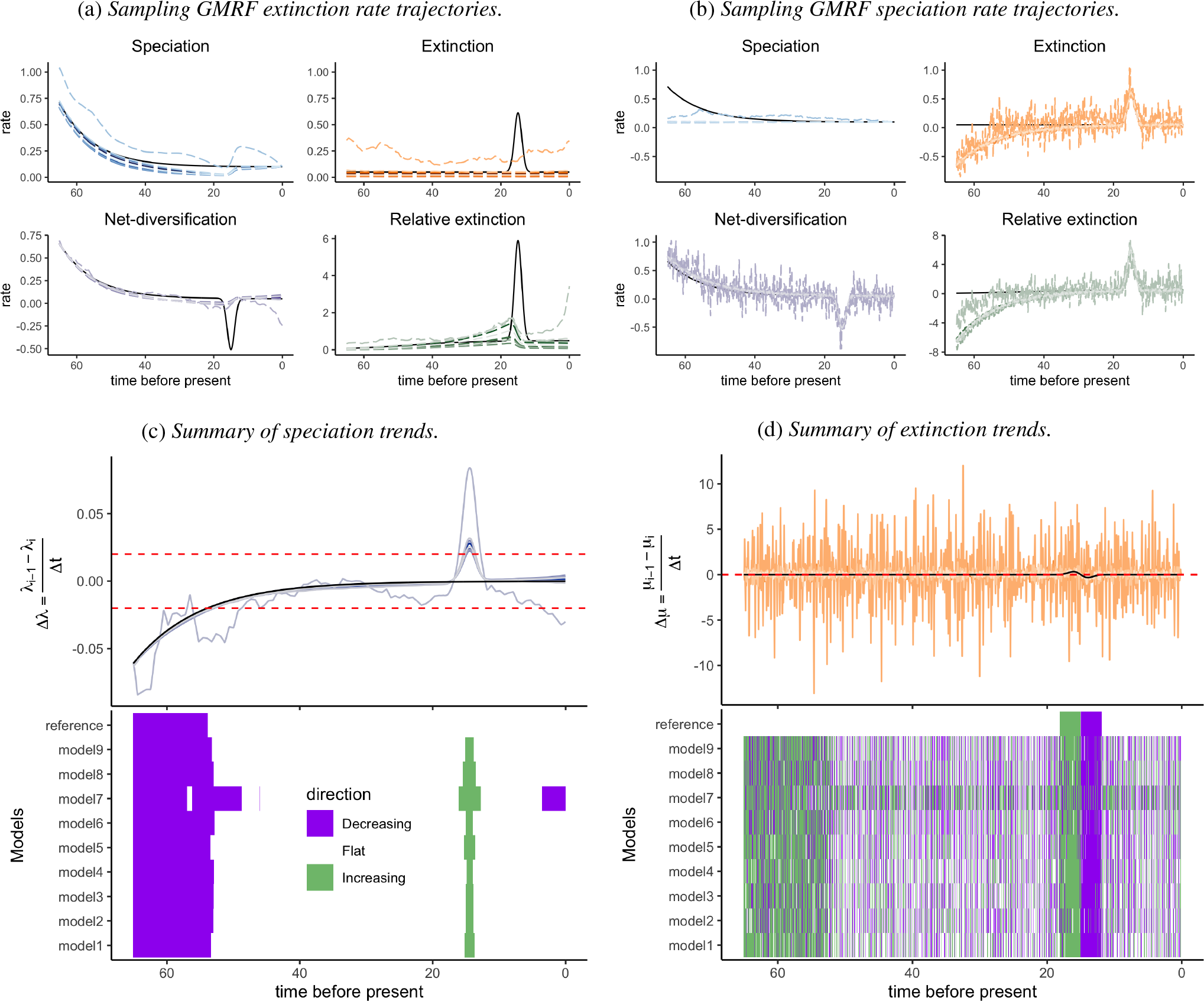
Illustration of diversification scenarios recovered with current congruence class sampling methods. Same as Figure 2, with speciation or extinction rates sampled as GMRF processes.

### A.3 Joint sampling with *p*-trajectories

**Figure A.3:**
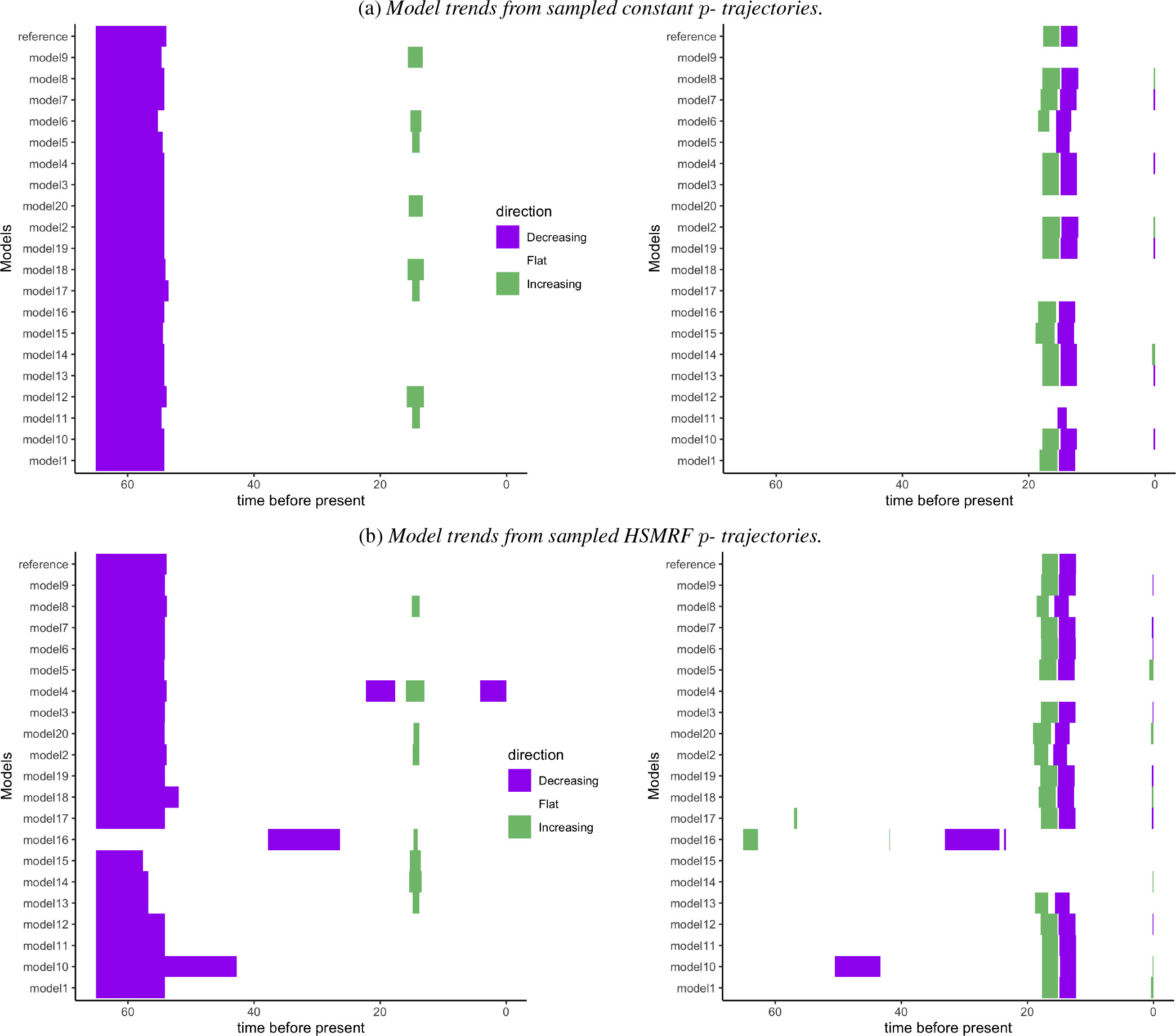
Summaries of the congruent diversification scenarios shown in Figure 3. The left panel summarizes speciation trends and the right panel extinction trends, both with a 0.02 threshold for significant increases or decreases.

## Appendix B Mammalian diversification histories

**Figure B.1:**
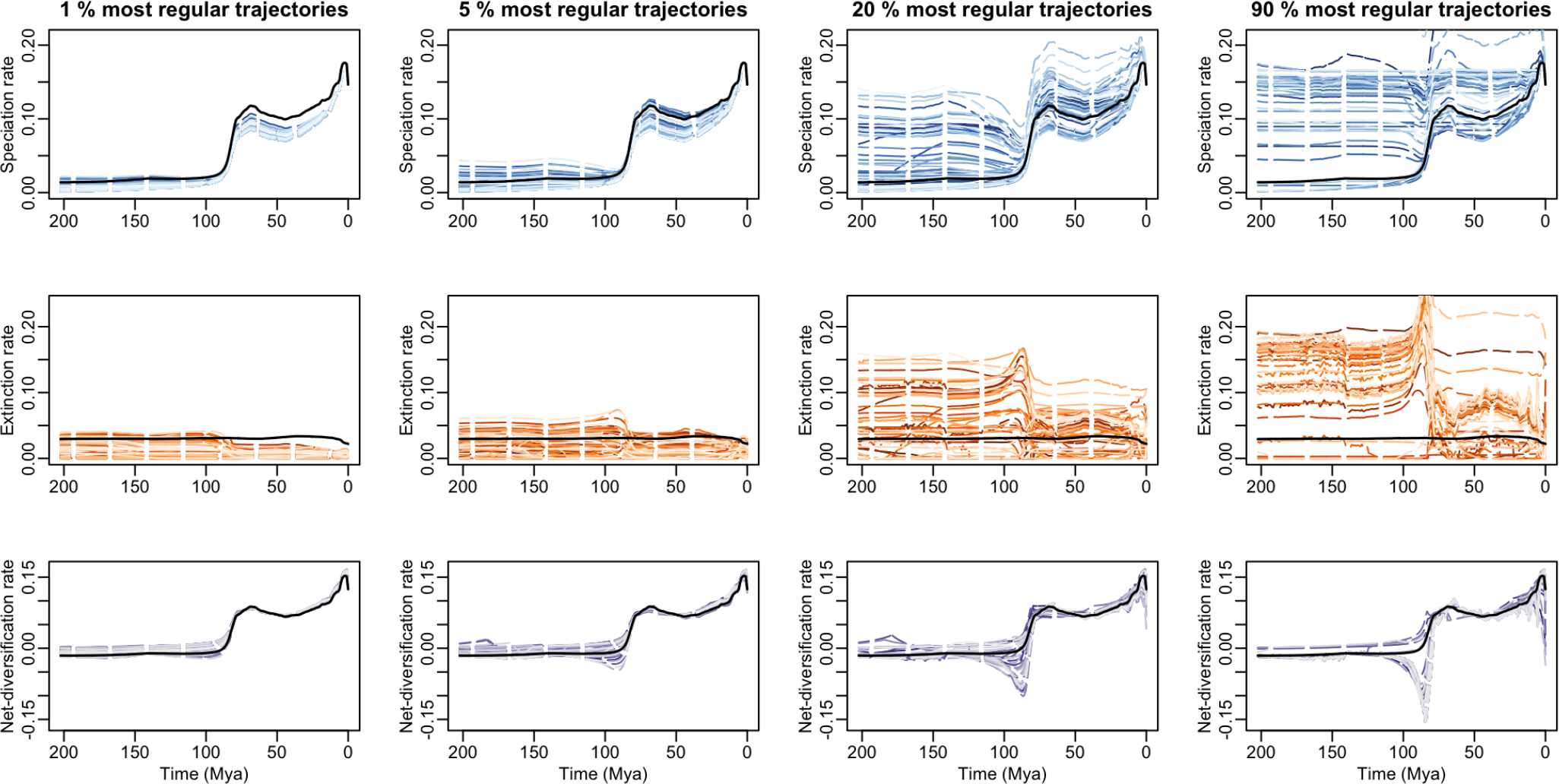
Congruent mammalian diversification trajectories for increasing regularity thresholds, using the ℒ 2 norm. 5000 joint HSRMF trajectories were sampled on the congruence curve, and then for each column 50 of them were selected among the most regular one (i.e. below the 1st, 5th, 20th, and 90th centiles, respectively). The chosen regularity criterion is a step-wise *ℒ* 2 norm on all the step-wise variations in both speciation and extinction rates.

**Figure B.2:**
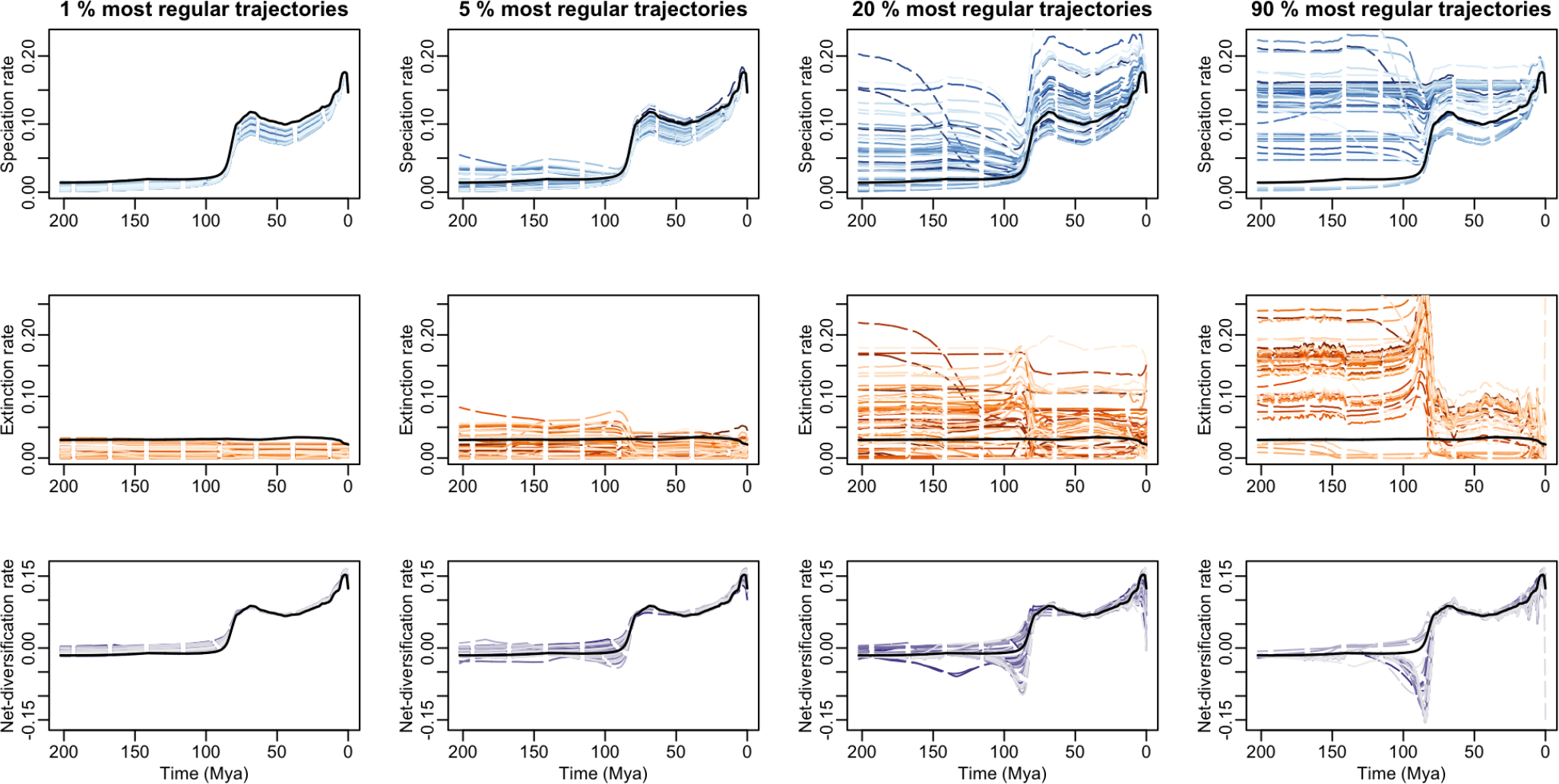
Congruent mammalian diversification trajectories for increasing regularity thresholds, using the ℒ 1 norm on derivative shifts. 5000 joint HSRMF trajectories were sampled on the congruence curve, and then for each column 50 of them were selected among the most regular one (i.e. below the 1st, 5th, 20th, and 90th centiles, respectively). The chosen regularity criterion is a step-wise **ℒ** 1 norm on all derivative shifts in both speciation and extinction rates.

**Figure B.3:**
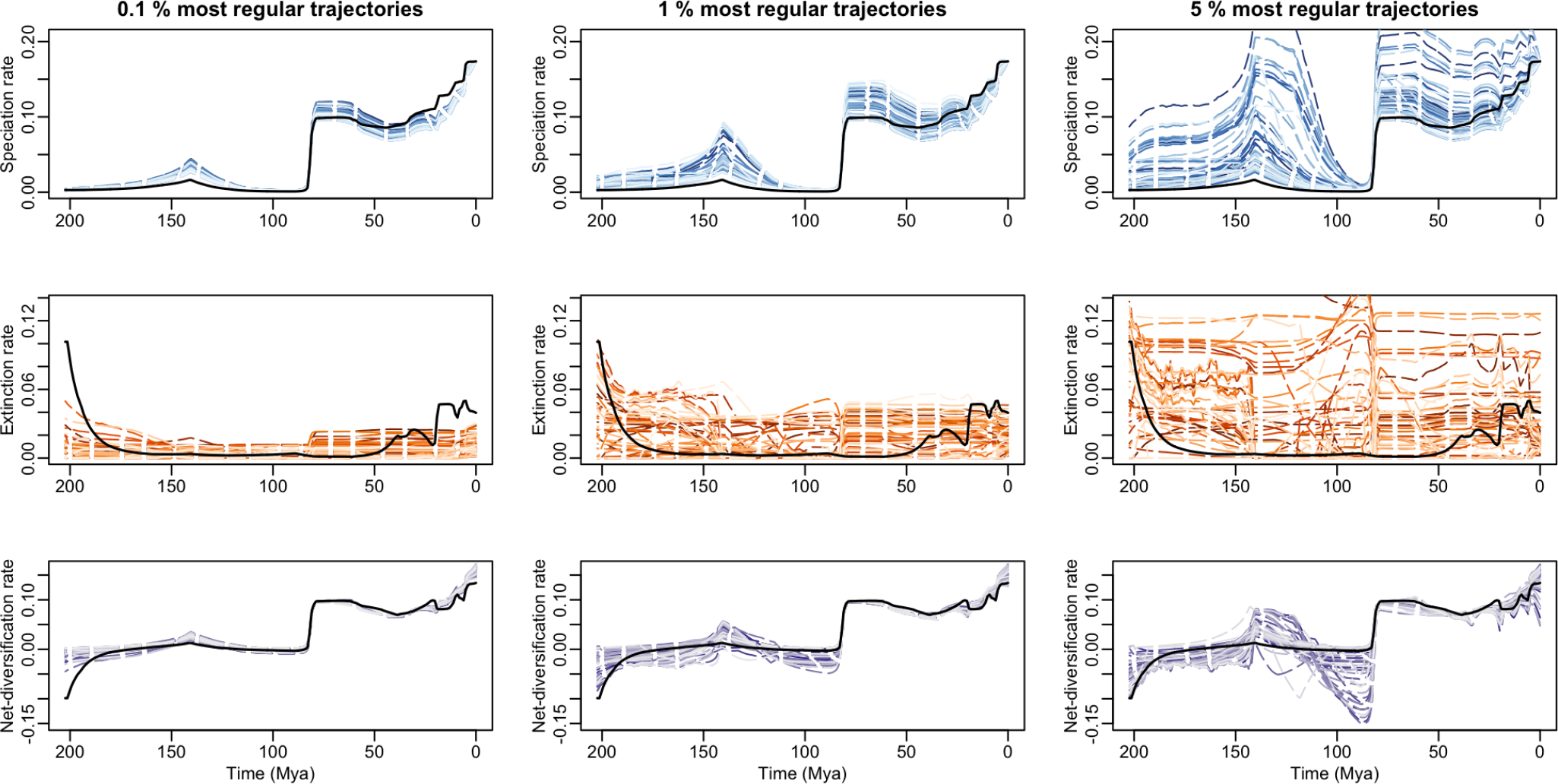
Congruent mammalian diversification trajectories for increasing regularity thresholds, without prior reguarization on rate shifts in the reference diversification analysis. The reference trajectory corresponds to the median rates from an estimation with an uncorrelated piecewise-constant model. 5000 joint HSRMF trajectories were sampled on the congruence curve, and then for each column 50 of them were selected among the most regular ones. The chosen regularity criterion is a step-wise **ℒ** 1 norm on all the step-wise variations in both speciation and extinction rates.

## Footnotes

1 https://revbayes.github.io/tutorials/divrate/ebd.html

